# IntegrIBS: Evaluating the Potential and Challenges of Building a Robust IBS Classifier with Integrated Microbiome Data

**DOI:** 10.1101/2024.06.21.600147

**Authors:** Dinakaran Palani, Nishad Bapatdhar, Renuka Manchiraju, Bipin Pradeep Kumar, Samik Ghosh, Sucheendra K. Palaniappan

## Abstract

Irritable Bowel Syndrome (IBS) is a condition that is quite complicated and shares its symptoms with other related diseases, making it difficult to diagnose. In this study, we initially trained machine learning models on individual microbiome datasets and tested their performance on other datasets, observing variability and low precision among them. To mitigate this, we hypothesised that integrating multiple publicly available microbiome datasets will capture a wide spectrum of microbiome variations across different geographies and demographics. Utilizing this integrated dataset, the XGBoost model achieved a mean accuracy of 0.75 with a standard deviation of 0.04 in 10-fold cross-validation, demonstrating its potential for robust IBS prediction. Explainability analysis identified key bacterial taxa influencing predictions, aligning with existing literature. However, the model’s performance declined significantly when using a leave-one-dataset-out approach, where the model was trained on all but one dataset and tested on the excluded dataset. The results highlight the challenges of generalizing across diverse datasets due to biological and technical variability. These findings present a cautionary tale regarding the integration of datasets and interpretation of results, emphasizing the need for more comprehensive approaches to develop reliable diagnostic tools for IBS.

## 1 Introduction

Irritable Bowel Syndrome (IBS) is a common gastrointestinal disorder which is often characterized by chronic abdominal pain, flatulence, bloating, and altered bowel habits for a set or a defined period of time. It is a complex and heterogeneous condition, with overlapping symptoms with other gastrointestinal disorders and its pathophysiology is not fully understood. Due to this, diagnosing IBS is complex. Very often, diagnosing IBS is done by the principal of exclusion of other conditions. Traditional diagnostic methods miss the intricate interactions within the gut microbiome that have been increasingly understood to contribute to IBS [1]. Collins [2] showed the influence of gut microbiota on gastrointestinal function and its potential as a therapeutic target for IBS. Andrews et al. [3] stressed the importance of managing the microbiome in clinical practice for IBS patients, suggesting personalized treatment strategies based on microbiome profiling.

An ideal diagnostic tool for IBS based on microbiome profiling would involve asking a patient with symptoms resembling IBS to undergo a microbiome profile of the gut through a stool test. Once the microbial abundances are known, an algorithm would analyze these abundances to ascertain if the person has IBS and help the clinician start the treatment regimen. To this end, we focus on developing a machine learning classifier to robustly predict IBS using microbiome data. To add, the microbiome composition can vary significantly due to demographic, environmental, and societal factors. To address this variability, it is crucial to integrate multiple datasets from diverse conditions to identify the core set of causative microbes. Thus, we integrated multiple publicly available datasets to enhance model robustness. Ford et al. [4] discussed the need for comprehensive datasets to better understand IBS, which aligns with our approach.

Previous studies using microbiome data for IBS diagnosis have shown potential. Camilleri [5] reviewed diagnostic and treatment strategies for IBS, noting the potential of microbiome-based approaches. Wang and Liu [6] compared different classifiers for human microbiome data, showing the effectiveness of machine learning techniques. Fukui et al. [7] demonstrated the utility of machine learning-based gut microbiome analysis in identifying IBS patients. Tanaka et al. [8] also used a machine-learning approach to diagnose IBS subtypes, showing the utility of advanced algorithms. However, these studies often relied on single datasets, limiting their generalizability.

Guided by these insights, our work started by integrating diverse datasets to capture a wide range of microbiome variations, in addition to performing dataset-specific processing steps which take into account nuances of data collection protocols of each study. This context specific processing and focus on being able to unify data cleanly could help in enhancing the predictive accuracy and generalizability of our IBS classifier. This aligns with Carcy et al. [9], who conducted a large-scale meta-analysis of IBS, highlighting the importance of comprehensive data integration. Using the integrated dataset, we trained an XGBoost model that achieved a mean accuracy of 0.75 with a standard deviation of 0.04 in 10-fold cross-validation, demonstrating the robustness of the integrated data classifier. Further, explainability analysis using SHAP identified 19 key bacterial taxa that influenced predictions that aligned well with existing literature.

However, the model’s performance declined significantly in the leave-one-dataset-out approach, where the model was trained on all but one dataset and tested on the excluded dataset. This discrepancy highlights critical concerns, potentially including over-fitting to the combined training data, dataset-specific biases, and insufficient generalization due to biological and technical variability across different datasets. These findings suggest that only a 16s based classifier may not be feasible given the current available datasets, and there is a need for more comprehensive signals, larger diverse datasets and multi-omic approaches to develop a robust and generalizable diagnostic tools for IBS.

The paper is organized as follows: Section 2 describes the datasets and preprocessing methods. Section 3 details the bioinformatics analyses and dataset-specific processing, outlining the pipeline and framework for building the IBS classifier. Section 4 presents the results, demonstrating the performance of combined datasets in the 10 fold cross validation study. We also provide, in Section 5, an explainable AI framework to understand the model’s predictions beyond a black box, which can further help in understanding the reasons for the model’s predictions. In Section 6, we train the classifier in a a leave-one-dataset-out approach, which revealed significant performance drops and highlighted the inherent challenges in generalizing across diverse datasets. This section highlights the need for caution when interpreting model performance in the context of heterogeneous 16s based microbiome data for IBS prediction. Finally, Section 7 concludes the paper and suggests future research directions.

## 2 Data curation and pre-processing

A thorough search of the National Center for Biotechnology Information (NCBI) databases was conducted to identify studies related to Irritable Bowel Syndrome (IBS) and the gut microbiome. Identified studies were downloaded; however, many lacked essential information such as primer details and metadata necessary to distinguish between IBS and healthy samples.

Uploaded data ideally should be raw data, it is often altered by various sequence trimming and merging techniques before submission. Many studies have not documented the alterations or trimming and merging techniques applied to their sequencer output. As a result, each study must be evaluated individually based on the quality of its data.

For this study, we began with 50 studies obtained from the NCBI website. Among these, 18 studies lacked associated literature but contained sequences and metadata, which were included in the analysis. Subsequently, of the remaining 32 studies, only 25 had metadata capable of distinguishing between IBS and control samples. Further refinement based on primer information reduced the selection to 19 relevant studies, after excluding those related to Internal Transcribed Spacer (ITS) and Whole Metagenome Shotgun Sequencing (WMGS). This process underscores the significance of documenting metadata and primer information in clinical trial research. The process is described in Fig. 1

**Fig. 1.**
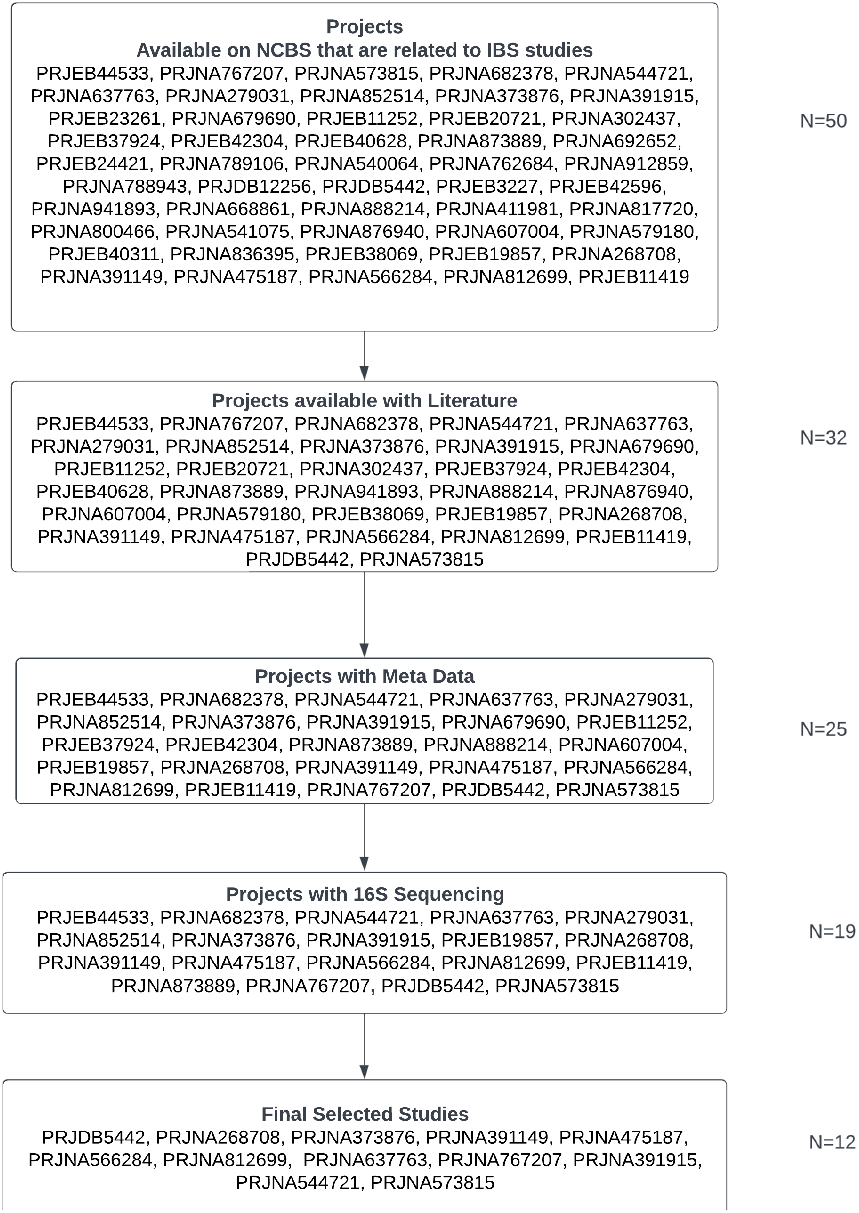
Selection Criteria and Process for IBS Project Dataset Inclusion

**Fig. 2.**
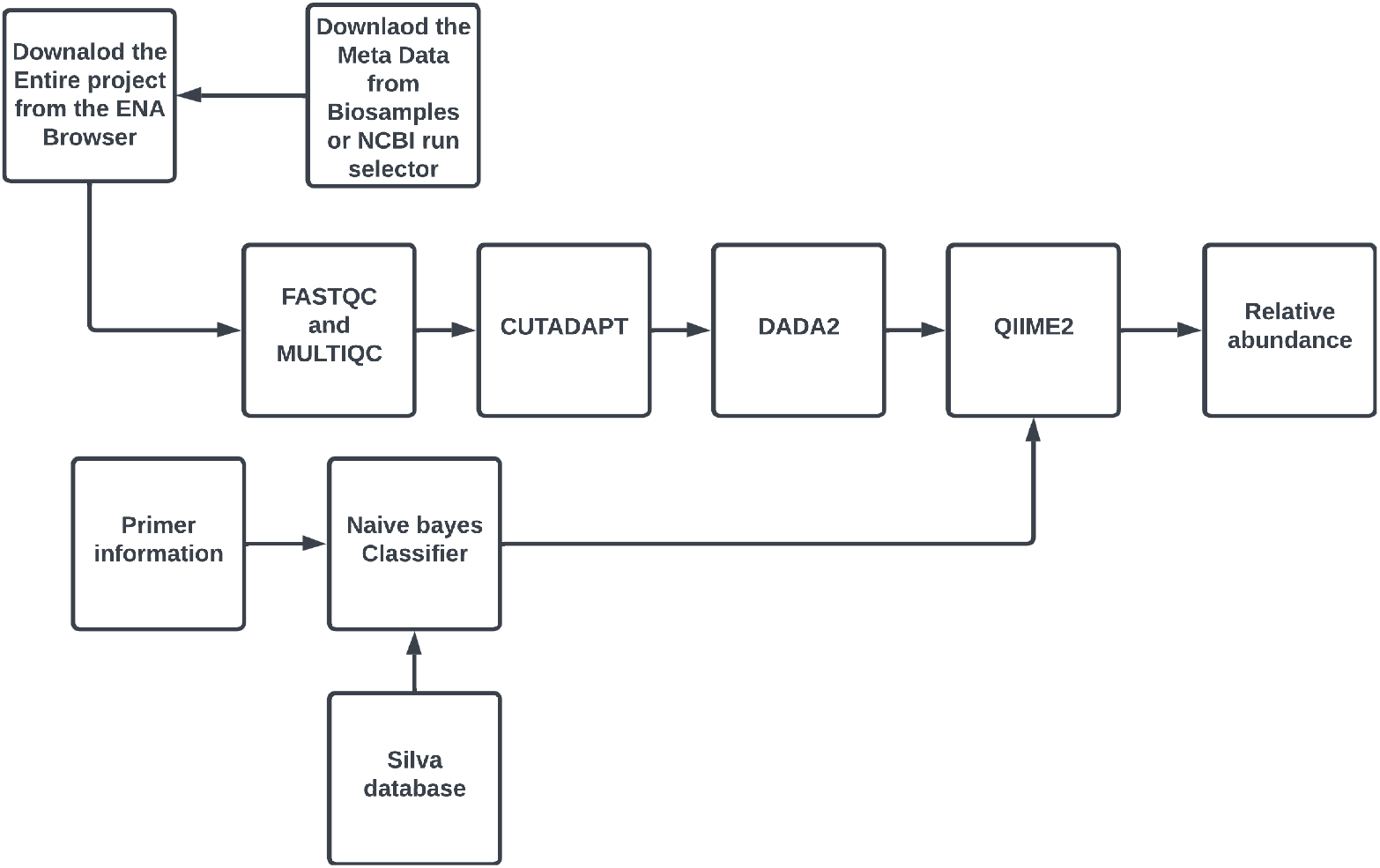
Block Diagram of the Bioinformatics Pipeline for 16S rRNA Sequencing Data Processing

Our analysis focused on gut microbiome data from the filtered 19 independent studies worldwide. Further, four studies were excluded due to challenges with *FastQC* and *DADA*2 resulting in insufficient reads, boiling down in a final selection of 12 studies. At the author’s request, one study data was made available[10]. Finally, these studies encompassed 1002 samples from individuals diagnosed with Irritable Bowel Syndrome (IBS) and 450 healthy control samples, with sample sizes per study ranging from 10 to

300. All datasets were downloaded from the European Nucleotide Archive (ENA)[11], and accompanying metadata were retrieved from the BioSamples database[12]. Table 1 summarizes the selected studies for this, listing their PubMed identifier (PMID), number of IBS and healthy control (HC) samples, data provenance, targeted regions, and the country of origin. This provides a clear snapshot of the dataset diversity and methodological details.

**Table 1.** Table of Final Studies Picked for Data Integration: Metadata and Provenance Details.

| PMID | # IBS | # HC | Data Provenance | Regions | Country |
| --- | --- | --- | --- | --- | --- |
| 28628918[13] | 10 | 10 | PRJDB5442 | V1-V2 | Japan |
| 26239401[14] | 202 | 88 | PRJNA268708 | V4 | Spain |
| 28457228[15] | 29 | 23 | PRJNA373876 | V3-V5 | USA |
| 33925672[16] | 36 | 41 | PRJNA391149 | V1-V3 | Italy |
| 30090090[17] | 17 | 9 | PRJNA475187 | V3-V4 | China |
| 31681624[18] | 15 | 14 | PRJNA566284 | V4 | China |
| 36624530[19] | 305 | 175 | PRJNA812699 | V4 | USA |
| 22180058[10] | 37 | 20 | PMID_22180058 | V4 | Sweden |
| 32727141[7] | 85 | 26 | PRJNA637763 | V1-V2 | Japan |
| 35704960[20] | 60 | - | PRJNA767207 | V3-V4 | Russia |
| 29708822[21] | 68 | - | PRJNA391915 | V6-V7 | USA |
| 32552668[22] | 84 | 44 | PRJNA544721 | V3-V4 | China |
| 33168839[23] | 54 | - | PRJNA573815 | V3-V4 | China |

### 2.1 Combining Datasets for building a IBS classifier

Despite having identical clinical objectives, individual studies can yield contradicting results regarding the same bacteria due to smaller sample sizes. This variability underscores the need for a larger, more diverse dataset to identify robust correlations between the microbiome and disease conditions. Additionally, gut microbiome diversity varies significantly by geographic location, suggesting that combining datasets from different regions can capture a broader spectrum of variations. Previous research suggests that microbial associations differ across populations, affecting the robustness and generalizability of microbiome studies [24, 25]. Our intial results also show that classifiers trained on single-study data perform poorly when tested on other datasets, highlighting the limitations of models developed from isolated datasets.

By integrating microbial profiles from multiple studies, we aim to leverage the increased sample size and diversity to develop a more robust microbial signature. We hopes that this integration might mitigate the limitations of small sample sizes and study-specific biases. However, integrating these datasets can also introduces new challenges, such as biological and technical variability, need for much larger sample sizes to account for the increased complexity and unknown variables inherent in a combined datasets. Our initial hypothesis was that integrating multiple datasets will enhance the predictive accuracy and robustness of our IBS classifier. To test this hypothesis, we compare the performance of machine learning classifiers trained on the combined dataset against those trained on individual datasets.

## 3 Data specific analysis pipeline

This section outlines the preprocessing and analysis pipeline used to address the dataset-specific nuances of 16S rRNA sequencing data from the studies we identified in the previous section. Our goal is to integrate diverse datasets of IBS vs healthy. Key steps in the pipeline include sequencing quality control, read trimming, inference of amplicon sequence variants (ASVs), training the classifier based on primer sequences, and taxonomical classification. Each step is structured to standardize and optimize the data for reliable downstream analysis.

- **Sequencing Quality Control:** Quality control of the raw sequencing reads was performed using FastQC and MultiQC. FastQC provides comprehensive statistics and visualizations to assess the quality of the sequencing data, while MultiQC aggregates results from multiple tools to provide a summary report. Raw FASTQ files were analyzed for quality scores across all bases, GC content, sequence duplication levels, and the presence of adapter sequences. Datasets may be discarded due to incomplete information causing failures in quality control analyses. The output reports were manually inspected to ensure the data quality was sufficient for downstream analysis.
- **Trimming of Reads:** Adapter trimming and quality filtering of the raw reads were conducted using Cutadapt (version 3.4) [26]. Adapter sequences were identified and specified for removal. Reads were trimmed to remove low-quality bases, ensuring a minimum quality. Post trimming, the quality of the reads was reassessed using FastQC to confirm the removal of adapters and improvement in read quality.
- **Infer Amplicon Sequence Variants (ASVs):** Amplicon sequence variants (ASVs) were inferred using the *DADA*2 pipeline [27]. The workflow involved multiple steps:
  – Filtering and Trimming: Reads were filtered to remove low-quality regions and trimmed to a uniform length to reduce error rates.
  – Error Learning: Error rates were estimated from the data.
  – Denoising: Sequences were denoised to infer true biological sequences, correcting for sequencing errors.
  – Chimera Removal: Chimeric sequences were identified and removed.
- **Training a Classifier Based on Primer Sequence for Sequence Classification of Bacteria:** A classifier was trained using a reference database tailored to the specific primer sequences used in this study. The reference sequences corresponding to the target region were extracted and formatted to train the classifier. This step ensured accurate taxonomic classification by aligning the reference database with the amplified region defined by the primers.
- **Taxonomical Classification:** Taxonomical classification of the inferred ASVs was performed using QIIME2’s feature classifier plugin method with the trained reference database. The classifier was trained on the SILVA database[28], with reference reads extracted based on the primer sequences used in the studies.

### 3.1 Pipeline Implementation

The curated datasets were processed using the nf-core/ampliseq pipeline, facilitated by the NF Tower platform. The nf-core/ampliseq pipeline is a robust and standardized workflow for processing amplicon sequencing data, particularly 16S rRNA gene sequences. The Azure Blob Storage and Azure Batch services were integrated to run the nf-core/ampliseq pipeline efficiently. Each study thoroughly inspected the generated MultiQC reports, and appropriate quality cutoffs were determined to minimize data loss.

### 3.2 Trimming Strategy Considerations

Trimming involves the removal of low-quality bases from sequencing reads, which can significantly impact the accuracy of downstream analyses such as alignment and assembly. Two primary approaches to trimming include cutoff lengths and quality-based trimming that are used by the DADA2 pipeline

- **Cutoff lengths (truncLen)**:
  – *truncLen* specifies the exact read length at which reads should be truncated. This is useful when there is a specific target read length or consistent read length. However, it may lead to the loss of valuable information if read lengths vary significantly or if there are substantial quality drops toward the end of the reads.
- **Quality-based trimming (***trunc*_*qmin*_, *trunc*_*qmax*_, *max*_*ee*_, *min*_*len*_):
  – *trunc*_*qmin*_ and *trunc*_*q*_*max* set the minimum and maximum acceptable quality scores for read truncation, respectively.
  – *max*_*ee*_ sets the maximum number of expected errors allowed in a truncated read.
  – *min*_*len*_ sets the minimum length a read must retain after truncation to be retained for further analysis. This approach allows for more flexible trimming based on quality scores, potentially retaining more data if quality allows.

### 3.3 Microbiome Dataset specific processing

The datasets were processed according to their unique characteristics. Below is a summary of each study and the specific pre-processing steps applied.

- **PRJNA573815**: This clinical study investigates the use of laxatives and probiotics for the treatment of Irritable Bowel Syndrome (IBS). The microbiome data is accessible via the ENA browser. Samples with the sample alias and baseline samples were selected for analysis. Primers were already removed, and merged reads were uploaded to the ENA browser. Multiple analyses were conducted with different cutoff lengths, starting from 445 and decreasing in increments of 5. A cutoff length of 425 was selected as it allowed a higher number of reads to pass through the DADA2 pipeline.
- **PRJNA544721**: This study aimed to identify the IBS phenotypes and healthy controls. Using the metatag isolation source, we identified samples of IBS-D and healthy controls. The primers were removed, and multiple analyses were conducted with cutoff lengths starting from 450 to 420. A cutoff length of 430 was selected because it allowed more reads to pass through.
- **PRJNA391915**: This study focuses on the effects of rifaximin on IBS-D patients. Baseline samples with read lengths of 150 and 140 were selected. Samples were split and run individually due to poor quality, and quality trimming parameters were applied. The final analysis included 68 samples, which were added to the database.
- **PRJNA767207**: This study examined different therapeutics for IBS patients. Samples were identified using metadata by looking at patient ID and visit number. Primers were removed, and a single read length was used. Analyses started with different cutoff lengths ranging from 455 to 415, with 420 chosen as the final read length.
- **PRJNA637763**: This study used machine learning techniques to classify patients as IBS or healthy. Samples were identified based on the description metatag. Double-ended reads with primers were removed using Cutadapt. Multiple combinations of reads were analyzed, with final cutoff lengths of 235 and 230 selected.
- **PMID-22180058**: This study defined IBS subtypes by species-specific alterations. Data access was granted via personal request, and demultiplexing was done using a provided mapping file. Single-ended reads with primers removed were processed with multiple Qiime2 classifiers.
- **PRJNA812699**: This study identified multi-omics data for IBS subtypes. Samples were differentiated using the gastrointestinal tract disorder tag. Primers were already cut, and cutoff lengths of 247 for forward reads and 237 for reverse reads were used.
- **PRJNA566284**: This study had two different cutoff length ranges, 219 and 270. Samples were batched based on read lengths, with a forward cutoff of 260 and reverse cutoff of 220 for the 270 batch, and 212 for both forward and reverse reads in the 219 batch. Primers were removed before analysis.
- **PRJNA475187**: This study aimed to find gut microbiome alterations in IBS-D patients. Samples were differentiated using the experiment-alias metatag. Primers were removed using Cutadapt, and cutoff lengths of 272 for forward reads and 247 for reverse reads were chosen.
- **PRJNA391149**: This pilot study identified dietary connections in IBS patients. Samples were classified using the metatag. Based on the percentage of input nonchimeric reads, samples were split into two groups. Primers were removed, and a cutoff length of 320 was used for single-ended samples.
- **PRJNA373876**: This correlation study examined the relationship between gut microbiome and brain volumes. Samples were identified using the host diseases tag, with values for IBS and healthy controls. Primers were removed, reads were merged into single-ended reads, and a cutoff length of 354 was used.
- **PRJNA268708**: This study observed reduced butyrate-producing bacteria in IBS patients. Samples were identified using the description tag to distinguish between IBS and healthy controls. Primers were removed, and a cutoff length of 310 was used.
- **PRJDB5442**: This study aimed to identify potential donors for fecal microbiota transplantation (FMT) in IBS patients. Samples were identified using the description tag. Primers were removed using Cutadapt, and a cutoff length of 320 was chosen.

### 3.4 From Taxonomical Classification to a building a Classifier for IBS

Once we get the taxonomic data from the above pipeline, next, these features are used to train a machine learning classifier that has the ability to distinguish between IBS patients and healthy controls. The process begins with the taxonomic profiles generated by QIIME2 or DADA2, which include relative abundances, presence/absence of taxa, and diversity metrics for each sample. These relative abundances are then used as features that serve as inputs for training machine learning models (classifier). Finally, the model predicts whether a given sample belongs to an IBS patient or a healthy control.

## 4 Results

In this section, we present our findings of our various experiments. First, we demonstrate that models trained on individual datasets perform poorly when tested across other datasets, highlighting the limitations of isolated dataset approaches. Next, we show that the results of training our best-performing model using the combined dataset Next, evaluated through 10-fold cross-validation which seemed to show promise with improvements in predictive accuracy and the model’s ability to pick the right features for IBS in explanability analysis was promising. Finally, we discuss the challenges observed when applying the leave-one-dataset-out approach, which reveals significant performance drops and underscores the complexities of achieving generalization across diverse datasets.

### 4.1 Classifier Training on individual datasets

Understanding the generalizability of classifiers trained on microbiome data from specific studies is important due to significant geographic variations in gut microbiome diversity. This experiment aimed to evaluate the performance of a classifier trained on one specific dataset when tested on all other datasets. It highlights the limitations of models trained on isolated datasets.

We implemented and compared four different machine learning models: XGBoost, Random Forest, a combination of XGBoost and Random Forest, and Naive Bayes. These models were chosen based on their performance in initial experiments and their use widely in the literature. For all the models we used model parameters as described in Table 3. In our initial experiments, XGBoost outperformed other models (as shown in the next subsection), prompting us to present the results for XGBoost here. Models were trained on individual datasets using taxonomic profiles generated by QIIME2 and tested across other datasets. The results are summarized in Table 2 for XGBoost (results for other models available on the supplementary website). The table shows the performance of XGBoost classifiers trained on one dataset and tested on others, including the number of healthy controls (HC) and IBS samples used for training and testing. For example, the first row of Table 2 shows the results for an XGBoost classifier trained on the dataset with PMID 33925672. This model was trained using 41 healthy control (HC) samples and 36 IBS samples. The trained model was then tested on a combined dataset consisting of 409 HC samples and 966 IBS samples from other studies.

**Table 2.** Performance of XGBoost Classifiers Trained on Individual Datasets and Tested on Other Datasets.

| Train PMID | Accuracy | Precision | Recall | F1 Score | AUROC | Train Data | Test Data |
| --- | --- | --- | --- | --- | --- | --- | --- |
| 33925672 | 0.544 | 0.737728 | 0.544513 | 0.626563 | 0.547019 | HC: 41,<br>IBS: 36 | HC: 409,<br>IBS: 966 |
| 36624530 | 0.48251 | 0.747449 | 0.420373 | 0.538108 | 0.557579 | HC: 175,<br>IBS: 305 | HC: 275,<br>IBS: 697 |
| 31681624 | 0.421644 | 0.786713 | 0.227964 | 0.353496 | 0.544028 | HC: 14,<br>IBS: 15 | HC: 436,<br>IBS: 987 |
| 30090090 | 0.593268 | 0.706844 | 0.702538 | 0.704684 | 0.53832 | HC: 9,<br>IBS: 17 | HC: 441,<br>IBS: 985 |
| 28628918 | 0.670391 | 0.707006 | 0.895161 | 0.790036 | 0.579504 | HC: 10,<br>IBS: 10 | HC: 440,<br>IBS: 992 |
| 28457228 | 0.469286 | 0.759009 | 0.346351 | 0.475653 | 0.58459 | HC: 23,<br>IBS: 29 | HC: 427,<br>IBS: 973 |
| 26239401 | 0.648881 | 0.701232 | 0.85375 | 0.770011 | 0.540824 | HC: 88,<br>IBS: 202 | HC: 362,<br>IBS: 800 |
| 22180058 | 0.635842 | 0.71577 | 0.785492 | 0.749012 | 0.556972 | HC: 20,<br>IBS: 37 | HC: 430,<br>IBS: 965 |
| 32552668 | 0.620091 | 0.706879 | 0.772331 | 0.738157 | 0.574442 | HC: 44,<br>IBS: 84 | HC: 406,<br>IBS: 918 |
| 32727141 | 0.630127 | 0.694367 | 0.820065 | 0.752 | 0.530703 | HC: 26,<br>IBS: 85 | HC: 424,<br>IBS: 917 |

**Table 3.** Model Parameters Overview. `n_jobs=8:` Utilizes all available CPU cores for computation, used in XGBoost and XGBoost+RF. `n_estimators=150:` Number of boosting rounds (in XGBoost) or trees (in Random Forest). `enable_categorical=True:` Supports categorical data directly in XGBoost. `random_state=1:` Sets a seed for the random number generator for reproducibility. `max_iter=1000:` Maximum iterations for solver convergence in Logistic Regression. Default parameters: Naive Bayes uses default settings.

| Model | Parameter |
| --- | --- |
| XGBoost | n_jobs=8, n_estimators=150, enable_categorical=True |
| XGBoost+RF | n_jobs=8, n_estimators=150, enable_categorical=True |
| Random Forest | n_estimators=150, random_state=1 |
| Logistic Regression | max_iter=1000, random_state=1 |
| Naive Bayes | Default parameters |

The results demonstrate that XGBoost classifiers trained on individual datasets perform inconsistently when tested on other datasets, reflecting the limitations of using isolated datasets for training. For instance, the classifier trained on PMID 33925672 achieved an accuracy of 0.544 when tested on the combined dataset of other studies, indicating poor generalizability. Similar trends were observed across other datasets, with accuracy and other performance metrics varying significantly.

### 4.2 Classifier Training on Combined Datasets with 10 fold cross validation

Next, We combined all available datasets into a single comprehensive dataset. This approach leverages the increased sample size and diversity, that could potentially address the limitations observed with individual datasets. To evaluate the performance and robustness of the combined dataset, we trained a model on the overall dataset and employed a 10-fold cross-validation (CV) strategy to give us a good estimate of the model’s generalizability.

XGBoost demonstrated the best performance, achieving a mean accuracy of 0.75 with a standard deviation of 0.04 in 10-fold cross-validation. Performance metrics included a precision of 0.79, recall of 0.88, F1 score of 0.83, and a ROC AUC of 0.81 (Table 4). The combined XGBoost and Random Forest model showed a mean accuracy of 0.73, with precision and recall scores of 0.77 and 0.88, respectively. Random Forest alone achieved similar accuracy (0.73) and a higher recall (0.94). Logistic Regression and Naive Bayes performed less effectively, with accuracies of 0.67 and 0.50, respectively (Table 4).

**Table 4.** Performance Metrics of Different Machine Learning Models on combined datasets.

| Model | Accuracy | Precision | Recall | F1 Score | ROC AUC |
| --- | --- | --- | --- | --- | --- |
| XGBoost | 0.75 ± 0.04 | 0.79 ± 0.04 | 0.88 ± 0.03 | 0.83 ± 0.03 | 0.81 |
| XGBoost+rf | 0.73 ± 0.03 | 0.77 ± 0.02 | 0.88 ± 0.04 | 0.82 ± 0.02 | 0.78 |
| Random Forest | 0.73 ± 0.03 | 0.74 ± 0.02 | 0.94 ± 0.03 | 0.83 ± 0.02 | 0.79 |
| Logistic Regression | 0.67 ± 0.04 | 0.72 ± 0.02 | 0.83 ± 0.05 | 0.77 ± 0.03 | 0.62 |
| Naive Bayes | 0.50 ± 0.04 | 0.81 ± 0.06 | 0.36 ± 0.05 | 0.50 ± 0.05 | 0.62 |

The ROC curves as shown in Fig 3 illustrate the performance of various classifiers in distinguishing between IBS and healthy samples. XGBoost achieved the highest AUC of 0.81, followed by the combined XGBoost+RF (0.78) and Random Forest (0.79). Logistic Regression and Naive Bayes showed lower AUCs of 0.62.

**Fig. 3.**
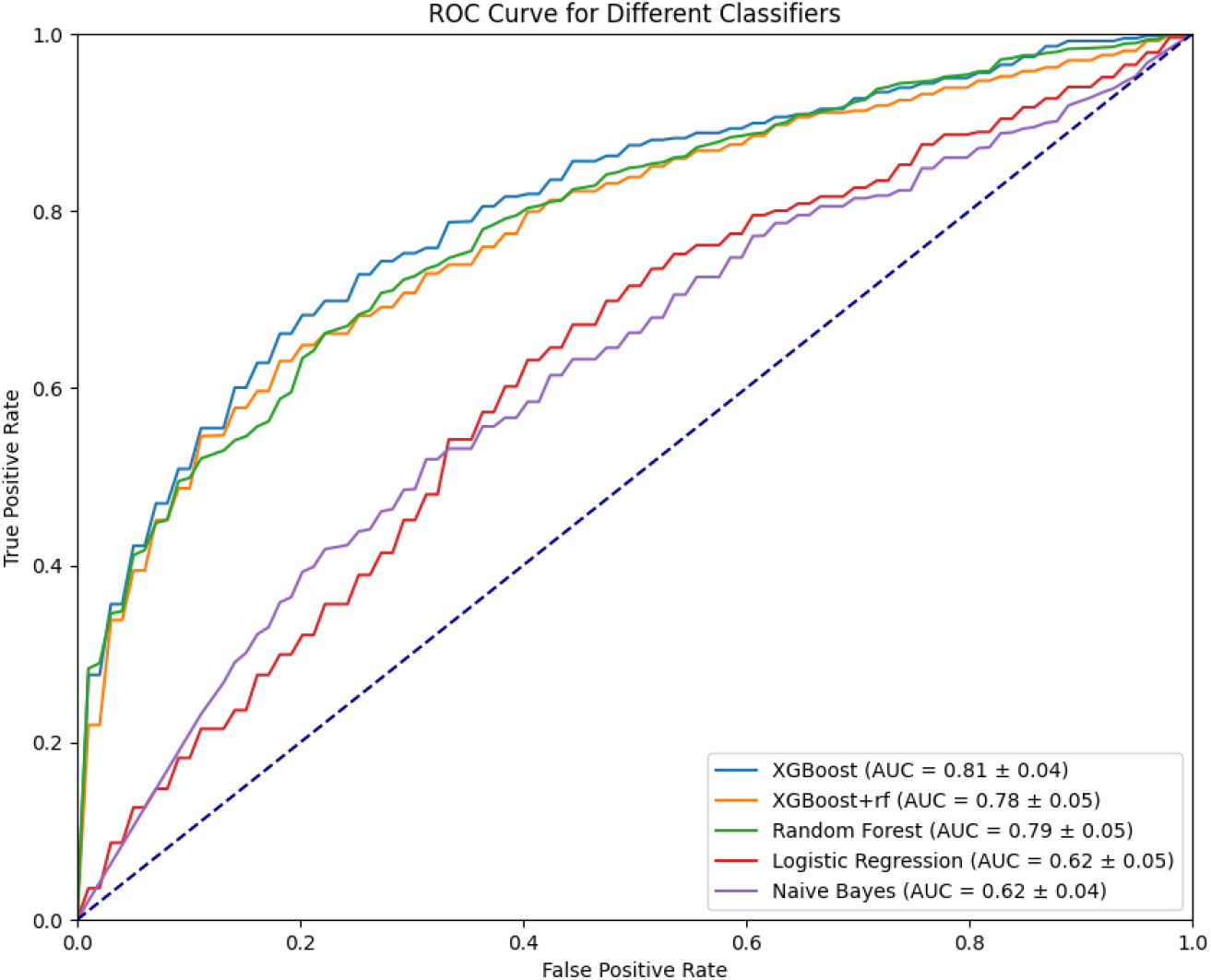
Receiver Operating Characteristic (ROC) Curves for Different Classifiers

### 4.3 Explainability Analysis of the integrated IBS Classifier Using SHAP

To gain insights into the model’s decision-making process, we employed SHAP (SHapley Additive exPlanations) values to interpret the XGBoost model. SHAP values help identify the contribution of each feature, in this case, specific bacteria, to the model’s predictions.

#### 4.3.1 Key Findings from SHAP Analysis

The SHAP analysis identified 19 key bacterial taxa influencing the IBS classifier’s predictions as shown in Fig 4. Notably, 16 of these bacteria have been previously mentioned in the literature in the context of their contribution to IBS. Table 5 summarizes the bacteria with positive and negative correlations to IBS, derived from SHAP values and supported by literature evidence. Positive correlations indicate that increased levels of these bacteria are associated with a higher likelihood of IBS, while negative correlations suggest the opposite.

**Table 5.**
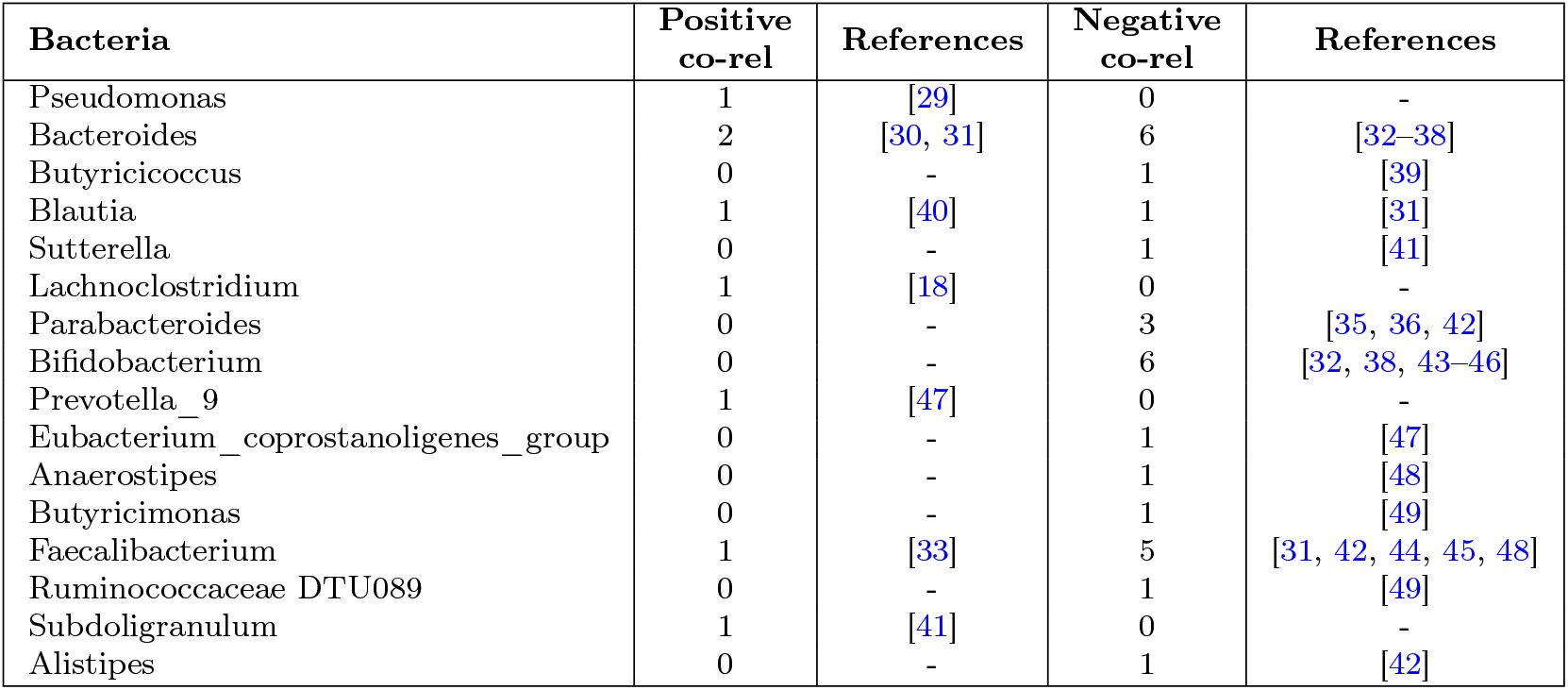
Summary of Bacterial Correlations with IBS: Positive and Negative Associations.

**Fig. 4.**
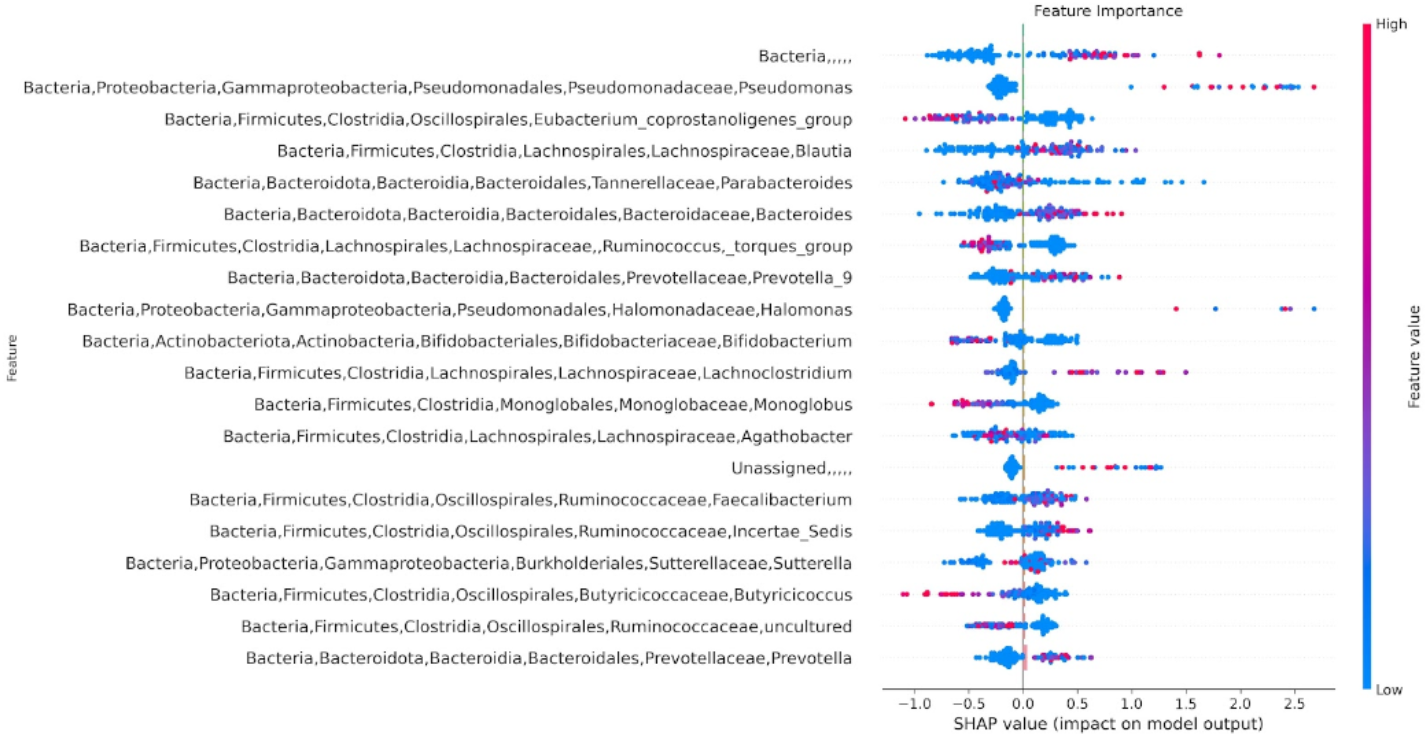
SHAP Values Indicating Key Bacterial Contributions to IBS Predictions by the XGBoost Classifier

### 4.4 Classifier Training on Combined Datasets with leave-one-dataset-out approach

Following encouraging results from the 10-fold cross-validation of the combined dataset, we next focused on evaluating the model’s robustness and generalizability. The 10-fold cross-validation experiments indicated improvements in accuracy and feature selection for IBS classification. However, to ensure the model’s applicability across diverse datasets and to simulate real-world scenarios where new, fully unseen data is encountered, we employed the leave-one-dataset-out approach. The leave-one-dataset-out approach essentially involves excluding one dataset for testing while using the remaining datasets for training the classifier. This process is repeated such that each dataset is used as the test set once. We wanted to check the model’s ability to generalize across different populations and study conditions, which could potentially rule out issues of over-fitting. For this experiment, we trained an XGBoost model. The pre-processing steps remain the same as before.

To our surprise, the leave-one-dataset-out approach showed significant performance drops compared to the 10-fold cross-validation results. The accuracies dropped significantly indicating that the models struggled to generalize when faced with entirely new datasets as shown in Table 6. Specific datasets such as PMID 33925672 highlight the inconsistencies in performance. Further, the classifier trained on the combined dataset but tested on PMID 33925672 showed a considerable decrease in performance, with an accuracy of 0.3636 and an AUROC of 0.3557, emphasizing the challenges of biological and technical variability between datasets.

**Table 6.**
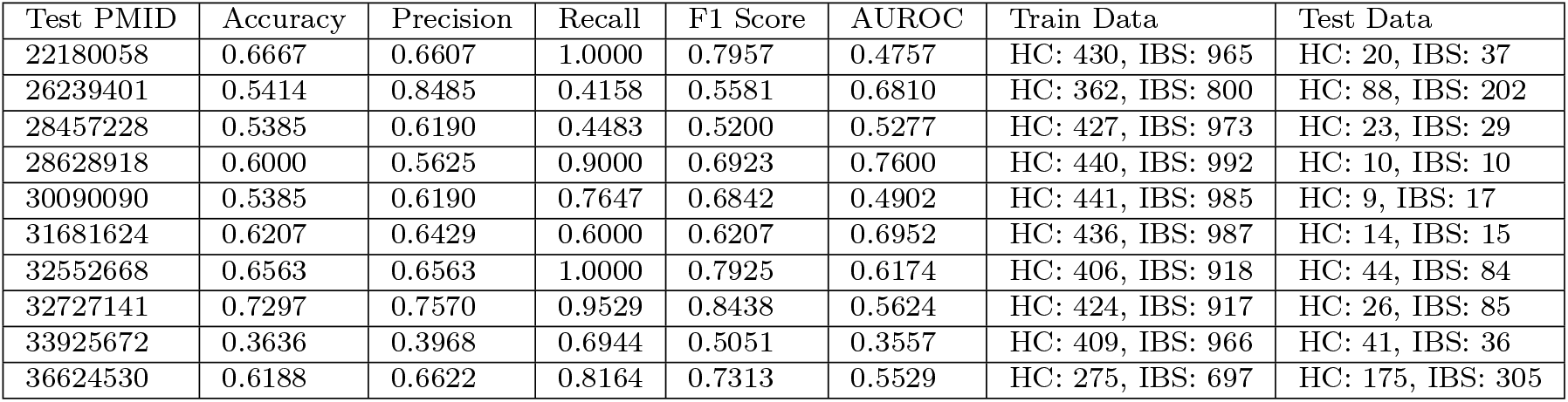
Summary of performance metrics for combined Datasets with leave-one-dataset-out approach.

| Test PMID | Accuracy | Precision | Recall | F1 Score | AUROC | Train Data | Test Data |
| --- | --- | --- | --- | --- | --- | --- | --- |
| 22180058 | 0.6667 | 0.6607 | 1.0000 | 0.7957 | 0.4757 | HC: 430, IBS: 965 | HC: 20, IBS: 37 |
| 26239401 | 0.5414 | 0.8485 | 0.4158 | 0.5581 | 0.6810 | HC: 362, IBS: 800 | HC: 88, IBS: 202 |
| 28457228 | 0.5385 | 0.6190 | 0.4483 | 0.5200 | 0.5277 | HC: 427, IBS: 973 | HC: 23, IBS: 29 |
| 28628918 | 0.6000 | 0.5625 | 0.9000 | 0.6923 | 0.7600 | HC: 440, IBS: 992 | HC: 10, IBS: 10 |
| 30090090 | 0.5385 | 0.6190 | 0.7647 | 0.6842 | 0.4902 | HC: 441, IBS: 985 | HC: 9, IBS: 17 |
| 31681624 | 0.6207 | 0.6429 | 0.6000 | 0.6207 | 0.6952 | HC: 436, IBS: 987 | HC: 14, IBS: 15 |
| 32552668 | 0.6563 | 0.6563 | 1.0000 | 0.7925 | 0.6174 | HC: 406, IBS: 918 | HC: 44, IBS: 84 |
| 32727141 | 0.7297 | 0.7570 | 0.9529 | 0.8438 | 0.5624 | HC: 424, IBS: 917 | HC: 26, IBS: 85 |
| 33925672 | 0.3636 | 0.3968 | 0.6944 | 0.5051 | 0.3557 | HC: 409, IBS: 966 | HC: 41, IBS: 36 |
| 36624530 | 0.6188 | 0.6622 | 0.8164 | 0.7313 | 0.5529 | HC: 275, IBS: 697 | HC: 175, IBS: 305 |

The performance degradation in the leave-one-dataset-out approach point to the complexities and challenges of generalizing models across diverse microbiome datasets for IBS. Several factors could be playing a role to this including differenes in microbiome composition due to geographic, dietary, and environmental factor, or challenges due to variations in sequencing techniques, sample processing etc (which we tried to tackle during our preprocessing steps) and over-fitting to the training data. For us the key take was was that while integrating multiple datasets improves predictive accuracy in a specific setting, it is not sufficient for achieving robust generalization. There is a clear need for more sophisticated approaches that can account for the inherent variability and complexity of microbiome data from 16s sequencing, including better sequencing technologies or using multi-omics approaches (data beyond just microbiome data to augment and strengthen the classification).

## 5 Discussions and Conclusion

In this paper, we aimed to demonstrate the potential benefits of integrating diverse datasets to improve the accuracy and robustness of an IBS classifier. Our initial experiments revealed significant variability and low precision when a classifier was trained solely on a single dataset and tested on others, highlighting the need for larger and more diverse datasets to enhance model generalizability. By combining several publicly available IBS datasets, which encompass different geographies and demographics, we attempted to account for a broader spectrum of microbiome variations, creating a comprehensive dataset for IBS classification.

In our 10-fold cross-validation on the combined dataset, models generally performed better, with XGBoost achieving the highest mean accuracy. SHAP analysis provided insights into the model’s decision-making, identifying key bacterial taxa influencing predictions. These identified bacteria aligned with existing literature, suggesting the biological relevance of the model.

Subsequently, we employed the leave-one-dataset-out approach for model performance evaluation, revealing significant performance drops compared to the 10-fold cross-validation results. This indicates that while the combined dataset approach showed promise in improving predictive accuracy within the cross-validation framework, it struggled to generalize to entirely new datasets. This variability points to inherent complexities arising from biological and technical differences across studies, such as variations in sequencing techniques, sample processing methods, and geographic diversity of microbiomes. Hence, in summary, we believe that merely merging datasets is not sufficient for developing a more robust IBS classifier.

Our findings resonate with those of Carcy et al.[9], who in their meta-analysis of IBS data, highlighted the importance of comprehensive data integration. They emphasized that while meta-analyses can identify robust microbial signatures, the integration of diverse datasets also brings challenges related to data heterogeneity and the need for advanced analytical methods to handle such complexity.

For future directions, multiple avenues can be explored. First, incorporating clinical parameters alongside microbiome data could provide a more holistic understanding of IBS and improve model predictions. Similarly, incorporating multi-omics data, including metagenomics, metabolomics, and transcriptomics, could offer a deeper understanding of the IBS-associated microbiome. Multi-omics integration can reveal interactions between different biological layers, offering a more comprehensive view of the disease. A key recommendation for the research community is to establish standardized protocols for sample collection, processing, and sequencing. This would help to reduce technical variability and improve the reproducibility of microbiome studies. Currently, the field of microbiome data analysis faces significant variability and inconsistency. Additionally, increasing the datasets with more samples from diverse populations will help capture the full spectrum of microbiome variability, thereby enhancing the generalizability of the models.

In summary, while our study presents a perspective on the potential of integrating diverse datasets for IBS classification, it also highlights the challenges and complexities involved. Addressing these challenges will be crucial for developing robust and generalizable diagnostic tools for IBS.

## Supplementary information

All data, code, models, and additional results of the paper are available online at https://bit.ly/IntegrIBS

## Acknowledgements

The Authors would like to acknowledge colleagues, especially Marcus Mata and Gobika Krishnan, for doing the literature review and downloading the databases. Author Sucheendra K. Palaniappan would like to acknowledge and thank the ONRG Grant for the Nobel Turing Challenge to The Systems Biology Institute (Grant number: N62909-21-1-2032) for funding support for part of this study.

